# Hybrid Raman spectroscopy and artificial neural network algorithm discriminating *mycobacterium bovis* BCG and members of the order *mycobacteriales*

**DOI:** 10.1101/2023.05.30.542797

**Authors:** Michael Macgregor-Fairlie, Paulo De Gomes, Daniel Weston, Jonathan James Stanley Rickard, Pola Goldberg Oppenheimer

## Abstract

Even in the face of the COVID-19 pandemic, Tuberculosis (TB) continues to be a major public health problem and the 2nd biggest infectious cause of death worldwide. There is, therefore, an urgent need to develop effective TB diagnostic methods, which are cheap, portable, sensitive and specific. Raman spectroscopy is a potential spectroscopic technique for this purpose, however, so far, research efforts have focused primarily on the characterisation of *Mycobacterium tuberculosis* and other Mycobacteria, neglecting bacteria within the microbiome and thus, failing to consider the bigger picture. It is paramount to characterise relevant Mycobacteriales and develop suitable analytical tools to discriminate them from each other. Herein, through the combined use of Raman spectroscopy and the self-optimising Kohonen index network and further multivariate tools, we have successfully undertaken the spectral analysis of *Mycobacterium bovis* BCG, *Corynebacterium glutamicum* and *Rhodoccocus erythropolis*. This has led to development of a useful tool set, which can readily discern spectral differences between these three closely related bacteria as well as generate a unique spectral barcode for each species. Further optimisation and refinement of the developed method will enable its application to other bacteria inhabiting the microbiome and ultimately lead to advanced diagnostic technologies, which can save many lives.

## Introduction

Despite the many advances in modern medicine, tuberculosis (TB) continues to affect millions of people every year (1). In 2021, the World Health Organization (WHO) estimated that active TB claimed 1.6 million lives (1). With the advent of SARS-CoV-2 and the resultant COVID-19 pandemic, it had been previously hypothesised that an increased number of TB patients could ‘fall through the cracks’ and would fail to acquire adequate treatment or diagnosis, especially in more rural areas (2, 3). The recent release of the Global TB Report 2022 has confirmed these hypotheses, with declining standards of treatment and diagnosis reported in most countries (1).

The primary causative agent of TB in humans is *Mycobacterium tuberculosis*. However, TB can be caused by other organisms within the *Mycobacterium* genus (4). This mainly consists of *Mycobacterium bovis* and *Mycobacterium africanum* (5), which together form part of the *Mycobacterium tuberculosis* bacterial complex (MTBC). TB infections primarily affect the respiratory system. In *circa* 90% of infected people, the bacteria enters the lungs and subsequently proceeds into a latent state (6). Whilst establishing an infection in this state, there is no propagation to such a degree that the bacteria can infect other individuals, nor do they exfiltrate the lungs. In effect, they remain in a state of relative equilibrium (7). However, in *ca*. 10% of patients, the bacteria progress onto causing an active disease (6). The patients start demonstrating symptoms such as fever, night sweats, weight loss and coughing, often with bloody sputum.

This spreads infection between individuals and can eventually prove fatal (8). In approximately 15% of people with an active infection the bacteria exfiltrate the lungs, causing extrapulmonary and systemic infection (9). Patients who are immunocompromised are more susceptible to this extrapulmonary disease (9). Eliminating TB infection requires an intense regime of antibiotics over the course of many months (10), which are often accompanied by an array of side effects including hepatotoxicity, peripheral neuropathy, gastrointestinal disturbances and arthralgia (11-13). Of a growing concern is the presence of antimicrobial resistance (AMR) within TB patients (14). This is particularly true for patients in the former USSR and India, which account for the highest burden of Multidrug-Resistant TB in the world (1, 15). Overall, the WHO seeks to reduce transmission of TB by ensuring both the rapid diagnosis of active cases as well as timely treatment of patients (16). Currently this presents with various issues due to the rural spread of cases. Diagnosis of TB often consists of molecular and immunological tests, which require specialist equipment and training often exacerbated by issues related to sensitivity and specificity or alternatively, requiring bacterial culture in centralised laboratories, which are slow and necessitate high containment facilities, rarely found in low and middle income countries where TB is endemic (17, 18). This has generated an increasing need for diagnostic techniques which are readily portable, fast, accurate, cheap, reliable and can be utilised at the point-of-need without specialist training (19).

Raman spectroscopy is a promising technique, which has previously been used to identify pathogens and various *Mycobacterium* sp. in monocultures (20) as well as in blood samples (21). In contrast to other diagnostic methods, Raman does not require an extensive sample preparation, addition of other reagents, complex laboratory equipment or specially trained personnel (22). Raman spectroscopy also renders itself easily miniaturised and portable and can be deployed outside the laboratory environment without compromising its performance. It is therefore, emerging as a potential candidate for rapid disease detection, which can be readily deployed in rural areas and address the unmet need in TB diagnostics (23). Raman spectroscopy has been successfully employed to discern between different types of bacteria (22), viruses (24) and several types of cancer (25). Spectral differences between various members of the *Mycobacterium* genus in monoculture have also been demonstrated (20). However, previous research focussed predominantly on a pre-requisite of culturing the bacteria, consuming valuable time whilst also requiring high-containment facilities (26). The use of blood samples as an indirect detection method was also considered (21). This has introduced further challenges in terms of requiring well-trained phlebotomists and difficulties with discerning TB within the smoking populations (due to the effects it has upon various serum proteins) along with a lack of consideration for HIV patients (21). Raman spectral signatures discerning *Mycobacterium sp*. from other bacteria comprising the microbiome within its full context or within real-world conditions have not yet been determined. Specifically, this could present issues relating to other bacteria from the order Mycobacteriales such as *Rhodococcus sp*. and *Corynebacterium sp*., which often form part of the microbiome (27), yet possessing similar structural elements to *M. tuberculosis* and other *Mycobacterium sp*. such as mycolic acid (28) and arabinogalactan (29, 30). It is, therefore, feasible that their Raman spectra may be similar in presentation.

Whilst research continues investigating Raman spectroscopy as a potential tool for TB diagnostics, it is imperative to establish a ‘baseline’ of the inherent characteristics of bacteria which compose the microbiome, where the detected changes *via* Raman spectroscopy could be attributed to the underpinning variations in bacterial physiology. The only study which has examined the related spectroscopic variations, is by Stöckel *et al*. (31) who used internal validation of samples *via* Leave-One-Batch-Out-Cross-Validation. While this model proved successful, no comparison of spectral changes due to other mycobacteriales outside the *Mycobacterium* genus were established to indicate spectral regions of change and the authors concluded that there is a need for advanced multivariate analysis due to the theorised high inter-variability between samples (31).

Here, we present Raman spectroscopy profiling and classification of *Mycobacterium bovis* BCG with a comparison of two other Mycobacteriales, in the hopes of establishing an important baseline in the form of a “multi-biochemical barcode”, as a characteristic tool for ongoing and future spectroscopic studies for diagnostic TB applications. We examine and evaluate what is the spectral variability between representative organisms present within the microbiome, using the three representative Mycobacteriales, *M. bovis* BCG, *Rhodoccocus erythropolis*, and *Corynebacterium glutamicum* and how these affect Raman spectra while gaining insights on the biochemical interpretation of spectral data.

The acquired spectral data is classified using our new artificial neural network algorithm, the self-optimising Kohonen index network (SKiNET) as a decision support tool. SKiNET is based on the separation of data classes in a self-organising map (SOM) with characterisation using a self-organising map discriminant index (SOMDI) enabling the subsequent classification of the tested data. Through inspection of key differences between neuron weights and class weight vectors, the algorithm enables identification of the key spectral changes. Training parameters used for the SOM included the grid size of 4, the learning rate of 0.5 and 10 epochs. From the separation of classes, it is evident that there are characteristic differences due to the obvious classification of certain neurons. As such, there is a clear basis for differentiation enabling characteristic weight vectors to be derived in the SOMDI. This data was then further analysed using PCA-LDA and then barcodes were generated. The identified barcodes from this study act as a reference, constituting a solid basis towards developing standard protocols as an essential prerequisite for reliable studies aimed at establishing the feasibility of Raman spectroscopy as an analytical tool for TB diagnostics. Molecular barcodes can further be constructed for distinguishing between TB-positive and TB-negative states associated with spectral changes *via* an easy subtraction of the variations from the reference sample spectra. In conjunction with the emergence of state-of-the-art machine learning techniques, the development of reliable and rapid spectroscopic analytical tools, this ultimately promises to improve diagnostic technologies, aiding in identifying a possible Raman based diagnostic technique for TB as well as a better quality and timeliness of disease diagnostics and tailored treatments.

## Materials and Methods

### Reagents

*Mycobacterium bovis* BCG (NCTC 5692) was cultured on Middlebrook 7H11 agar (ThermoFisher) + 0.2% Glycerol + 10% OADC Growth supplement (Sigma Aldrich) before being moved to Middlebrook 7H9 liquid media + 0.2% Glycerol + 10% OADC Growth supplement. *Corynebacterium glutamicum* (ATCC 21850) was cultured on Tryptic Soy Agar (Oxoid) prior to being moved to Tryptic Soy Broth (Oxoid) *Rhodococcus erythropolis* (ATCC 4277) was cultured on Brain-Heart Infusion Agar (Becton Dickinson) prior to being moved to Brain-Heart Infusion broth (Becton Dickinson).

### Sample Preparation

*M. bovis* BCG was cultured on agar for 4 weeks at 37°C on solid media. The colonies were subsequently placed in liquid broth, where they were incubated at 37°C and 150RPM for 1 week. *C. glutamicum* and *R. erythropolis* were cultured on the media at 37°C for 48 hours and then transferred to their respective liquid cultures and incubated for a further 48 hours at 37°C and 150 rpm. Following the incubation, cells were centrifuged at 3900 rpm for 10 mins and the supernatant was removed. The cells were then resuspended in 1ml of sterile, distilled water and spun again. This was repeated three times (20). Cells were aseptically removed from the respective pellet using a 10μl inoculating loop and spread on to an autoclaved aluminium coated glass slide. The cells were then allowed to air dry under laminar flow for two hours (20).

### Raman Spectroscopy

Spectral acquisitions were performed using an InVia confocal Raman (Renishaw). The spectrometer was calibrated prior to each use with silicon (520.7cm^-1^). A 100x Leica objective and a 1200l/mm grating were used for all measurements in the range of 750-1750cm^-1^. Raman map scans, consisting of 3 accumulations and 10 second exposure time, were acquired over a 10μm x 10μm square grid using a 785nm excitation laser with laser power of 10-14mW. 100 spectra were collected for each bacterium with 3 replicas, generating a total of *n*=300 spectra per bacterial species. Wire 5.1 (Renishaw Plc) software was used for baseline subtraction and cosmic ray removal.

### Principal Component Analysis

Principal component analysis (PCA) was used to interpret the *minute* differences in spectra between the different bacterial strains. PCA is a multivariate data analysis technique which reduces the dimensionality of the dataset into the most relevant components to maximize the variance among different samples by projecting the data into a space with N orthogonal basis vectors, arranged in descending order with respect to variance. Each axis, or principal component, represented a fraction of the total data variance. The dimensionality reduction was achieved by selecting the number of principal components that explains at least the required fraction of the dataset variance. In our study, this value was 99.9% of the total variance. PCA analysis was used to determine how the separated clusters separated between different strains and also to identify what they had in common. An ellipsoid was fitted to the cluster, which covers up to one sigma significance according to the respective data dispersion in the principal component one (PC1) and PC2. PCA loadings were calculated using the eigenvectors and eigenvalues obtained from matrix operations to find the PC. The loadings contained information specific to the initial Raman data and differed depending on the PC space. In a 2D PCA system, the loadings of the different PC quadrants were expressed as PC1 > 0 and PC2 > 0, PC1 0 and PC2, and PC1 > 0 and PC2 0, resulting in four different loading fingerprints related to the four quadrants.

### Principal Component Analysis and Linear Discriminant Analysis

Further, PCA was used as a pre-processing tool to reduce the dimensionality of the dataset and then used as input data to the linear discriminant analysis (LDA). The PCA pre-processing step yielded an inherent benefit of all of the output data being orthogonal, which circumvented a weakness of LDA *i*.*e*., the collinearity. LDA similar to the PCA, seek to maximise variance between components however, instead of within the dataset, LDA maximised variance between groups. This meant that it has projected the data into a space such that the variance between each bacterium was maximised. The comparison of the pure PCA and PCA-LDA approaches were compared for their relative effectiveness in classifying spectra and to the self-organising maps classification approach.

### Self-Organising Maps

The self-organising maps (SOMs) were used for multivariate analysis (32)-(33). SOMs are single-layer artificial neural networks that are represented as a two-dimensional (2D) hexagonal array of neurons. Inspired by the visual cortex in the brain, the SOM is trained to activate neighbouring neurons based on similar inputs, in this case Raman spectra. Each neuron has a weight vector with a length equal to the number of variables in a spectrum. The weights are gradually adjusted to be similar to the input data by exposing the network to training samples over several iterations, so that each neuron only activates on a specific spectral signature. The result is a 2D projection of hyperspectral data that can be seen as visible clustering based on type, group, and state. SOM employs the self-organizing map discriminant index (SOMDI), which appends a set of label vectors to each neuron and allowed us to study the most prominent features that cause the activation of a specific neuron to a class label. Following that, a supervised learning step was introduced to optimize the network and the class label associated with each neuron was used to quickly identify new data presented to the SOM.

### Raman Peak Analysis

The most important peaks were extracted by applying a multi-Lorentzian peak fitting method (33) followed by a peak comparison among the different samples and selecting the common peaks between samples to perform statistical analysis. A box plot of the Raman intensity of these peaks was performed and compared among samples using multiple pairwise comparisons, Tukey’s honestly significant difference test (Tukey’s HSD) to investigate the differences. Each spectrum was baseline subtracted and normalized between 0 and 1 using the asymmetric least square method.

## Code Availability

The customized written Python algorithm can be downloaded from Ref. (34).

## Results and discussion

Representative average Raman spectra in **Fig 1a** show the spectral differences between replicates and the bacteria analysed in this study with the variance of the sample highlighted with distinctive bands arising from the *R. Corynebacterium, M. Bovis* BCG and *R. erythropolis*. A higher sample variance can be seen between *R. erythropolis* and *M. bovis* BCG *via* the greater heterogeneity in the spectra. Clustering of the different bacteria obtained *via* the PCA (**Fig 1b**) with the greatest variability and therefore, the greatest cluster domain spread is observed in *R. erythropolis* and *M. bovis* BCG also evident from the Raman spectra (**Fig 1a**). The initial PCA analysis of the data also shows a great amount of spectral crossover.

**Fig 1.**
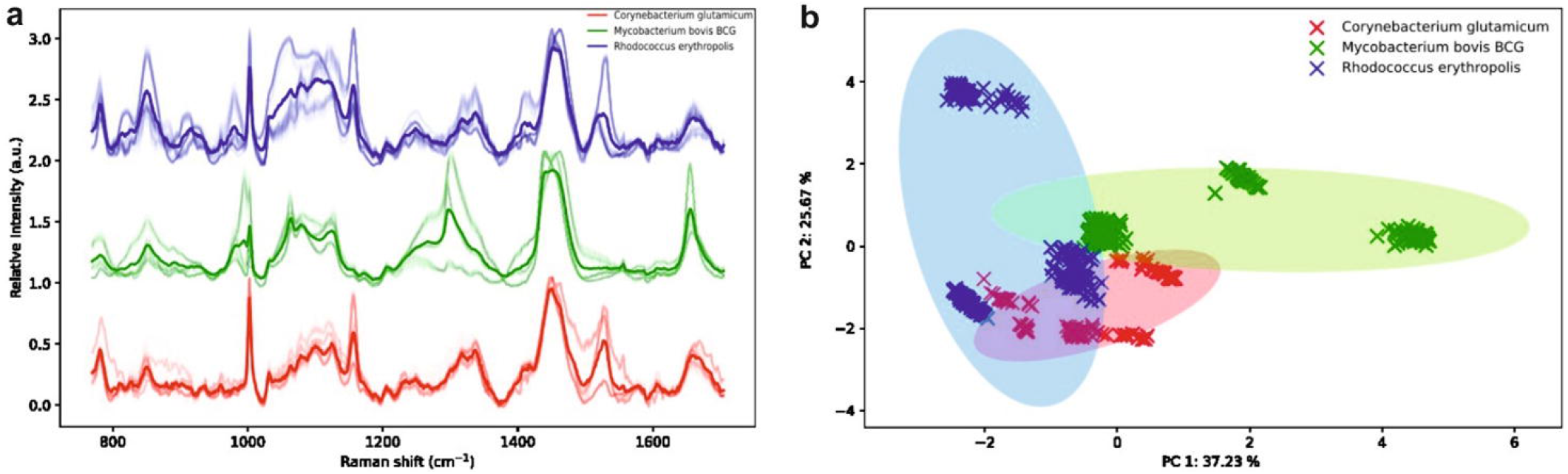
**(A)** Average Raman spectra of the various bacteria showing the variance within the same sample (faint lines) and the representative spectral fingerprint peaks (bold lines). **(B)** The corresponding PCA analysis of the bacteria.

Subsequently, to the PCA quadrant analysis of the bacteria samples, the loadings were analysed in terms of the specific quadrant location (**Fig 2a**). A high population of *R. erythropolis* and *C. glutamicum* samples in the 3rd, green quadrant (Q3) was observed with the corresponding loading spectra (**Fig 2b**) found to be related to the Q3 loading identified by the presence of multiple peaks, highlighting the relative similarities between both species of bacteria. *M. bovis* BCG located in Q1, shows a high signal intensity for the 1050cm^-1^ and 1300 cm^-1^ bands resulting in greater separation of the *M. bovis* BCG relative to the other bacteria. *C. glutamicum also* exhibited a cluster in Q4 due to the presence of a high-intensity peaks at 1150cm^-1^ and the 1300cm^-1^ region and the *R. erythropolis* shows a separated cluster in Q2, which is related to the presence of a high-intensity peak at 1050cm^-1^ combined with other peaks present in Q3 yet, with the absence of a strong peak at 1300cm^-1^.

**Fig 2.**
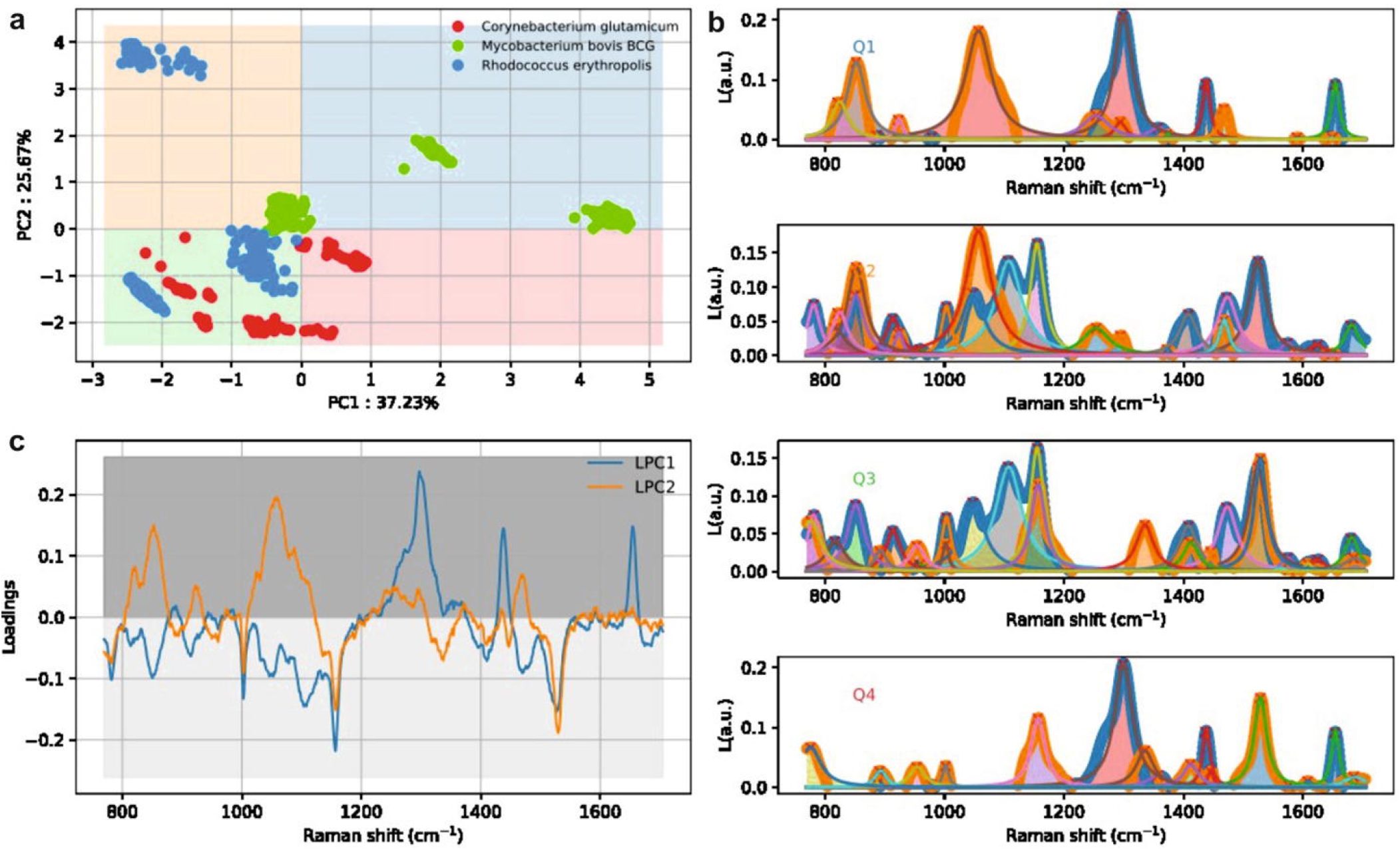
**(A)**. Quadrant analysis of the PCA data, illustrating the clustering in Q3. **(B)**. Clustering plot illustrating the most influential spectra in determining the clustering effects observed within the respective quadrants Q1, Q2, Q3, Q4. **(C)**. Raman loadings plot demonstrating the most influential parts of the spectra in the PCA.

Further, a region where most samples converge was identified between [-3,1] in PC1 and [-3,1] in PC2 thus, revealing a commonality among different samples although both *M. bovis* BCG and *R. erythropolis* have a larger variation in spectral signal which enables them to populate other regions of the PC space. This has also been identified from the Raman loadings plot (**Fig 2c**), where a few spectral lines above and below the sample average were indicative of the Raman signal variance.

This variance has further been identified *via* SOM (**Fig 3a**), where the sample separations clustering is identified as a colour intensity. The darker green hexagon is indicative of a more distinct spectral region for *M. bovis* BCG with the lighter green hexagon close to blue and the bright red hexagons, represent a degree of crossover between *M. bovis* BCG, *R. erythropolis* and *C. glutamicum*, respectively, further consolidating the PCA observations. *R. erythropolis* (blue) and *C. glutamicum* (red) appear mostly similar except the different peak intensities in the 1000cm^-1^-1200cm^-1^ region as well as the 850cm^-1^ and 1550cm^-1^ peaks and *M. bovis* BCG (green) appears to be more distinctive especially, in the 1200cm^-1^-1400cm^-1^ region along with an absence of a peak at 1550cm^-1^ and a more intense peak at 1700cm^-1^. The activation peaks (**Fig 3b**) correspond to the peak statistical analysis with *R. erythropolis* (blue) and *C*. glutamicum (red) being predominantly similar whilst *M. bovis* BCG (green) exhibiting is more distinct spectral fingerprint particularly, in the 1300cm^-1^ region.

**Fig 3.**
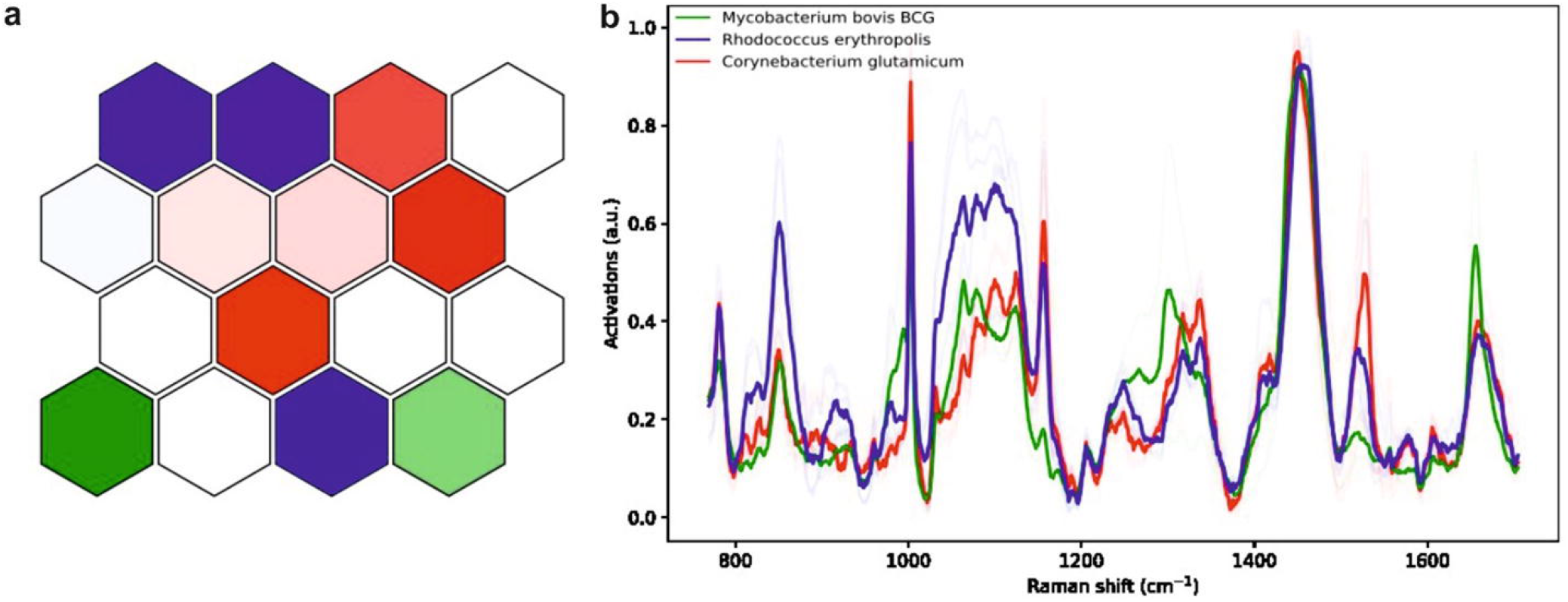
**(A)**. SOM trained to analyse the respective bacteria illustrating the clustering of the spectra. **(b)**. SOMDI showing the peaks which have the greatest influence in activating the respective neurons in SOM.

We have subsequently identified and statistically analysed the most dominant spectral peaks for the obtained Raman spectra (**Fig 4a**). Boxplots comparing the peaks shown in **Fig 4a** show the main differences between the peaks for each bacteria studied. The 780cm^-1^ peak is found to be similar for the *C. glutamicum* and *R. erythropolis* and significantly different for *M. bovis* BCG. This peak corresponds to an uracil containing ring, with the difference caused by the varying levels of uracil utilisation between the cells. Similar differences are also observed for 1155, 1450, 1525, and 1610cm^-1^ peaks. Highly significant peaks identified among all samples including the 1000, 1575 and 1660cm^-1^ are characteristic of Mycobacteriales. On the other hand, the peak at 1210 was found to be not statically significant. Peak differentiation can be achieved, mainly *via* the 850cm^-1^ band, which has been found to be statistically significant for the *R. erythropolis* (blue) but not for the *M. bovis* BCG (green) and *C. glutamicum* (red). We can, therefore, accurately identify the various bacterial species within the mixture. Specifically, *R. erythropolis* (blue) *via* the prominent intensity of the 850 cm^-1^ peak, associated with tyrosine, *C. glutamicum* (red) from the *M. bovis* BCG (green) *via* the 780, 1000, 1155, 1525 and 1610 cm^-1^ peaks, attributed to the uracil-based breathing ring, phenylalanine, carotenoids, in-plane vibrations of conjugated -C=C- and cytosine, respectively (**Table 1)**.

**Table 1:** Summary of the prominent peaks with the greatest statistical significance determined in **Figure 4A**. Peaks attributions are taken from Movasaghi *et* al. (39).

| Raman Peak ($\text{cm}^{-1}$) | Functional Group /<br>Band Assignment | Prominent In: | | |
| --- | --- | --- | --- | --- |
|  |  | *C.*<br>*Glutamicum* | *R.*<br>*Erythropolis* | *M. Bovis*<br>*BCG* |
| 780 | Uracil-based breathing ring | X | X |  |
| 850 | Tyrosine |  | X |  |
| 1000 | Phenylalanine | X | X | X |
| 1155 | Carotenoids | X | X |  |
| 1210 | Stretching in tyrosine and<br>Phenylalanine | X | X | X |
| 1450 | $\text{CH}_2$ bending | X | X | |
| 1525 | In-plane vibrations of conjugated -<br>$\text{C=C-}$ | X | X | |
| 1575 | DNA bases | X | X | X |
| 1610 | Cytosine | X | X |  |
| 1660 | Lipids | X | X | X |

**Fig 4.**
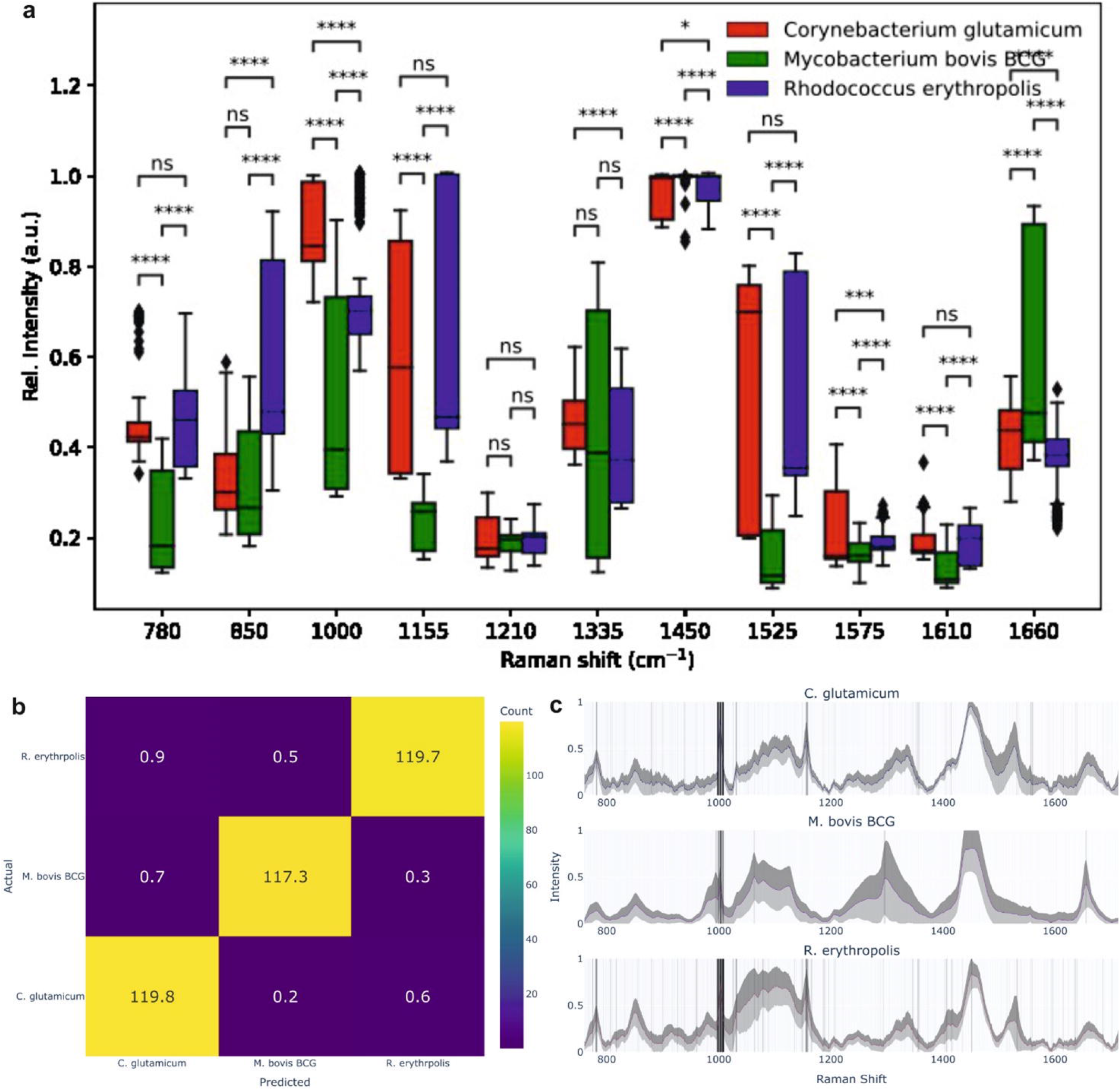
**(A)**. Box plots comparing the dominanat Raman spectral peaksof the studied bacterial spieces (NS = not significant, * = *p*<0.05, ** = *p*<0.01, ***= *p*<0.001, ****= *p*<0.0001). **(B)**. Confusion matrix following the 60:40 training and test data split during the PCA-LDA. **(C)**. Characteristic derived spectral barcodes of the *C. Glutamicum, M. Bovis* and *R. Erythropolis* after the application of a Savitzky-Golay filter and reduction of the noise from the double differentiation.

With the dataset pre-processed by PCA, the LDA data was subject to cross-validation, with 4 folds used in analysis. The mean accuracy (+1o-) was 99.11 + 0.63%, showing excellent discrimination between the different species (**Fig 4b**), where each species is clustered significantly apart from the rest with the overall detailed analysis partitioning the PCA-processed dataset into 40% test data and 60% training.

The derived confusion matrix (**Fig 4b** and **Table 1**) shows that on average the PCA-LDA approach is highly effective at discerning between different bacterium species. The off-diagonal values represent the incorrect identification occurrence, where on average each of these values was below 1. This means that in the worst performance case scenario, the PCA-LDA approach will misidentify two samples from a pool of three bacteria.

The obtained spectra were further classified to derive the characteristic fingerprint spectral barcode for each bacterium derived from the second derivatives, after application of a suitable threshold to discriminate between signal and noise (**Figure 4c**). Initially, a Savitzky-Golay filter was used to smooth the signal and reduce the noise from double differentiation and subsequently, a grid approach was used to select the window size and polynomial order with optimal values for these were identified as 5 and 2, respectively. Following filtering, the second derivatives were assigned binary values, +1 if their absolute value was greater than or equal to 5% of the maximum value for each species, and 0 otherwise. In our case, this threshold, building upon the work in Ref. (35), was increased to 5% since the 1% value applied in Ref. (35) unnecessarily included noise. If the value was below the threshold, it was assigned 0. Bars were plotted for each wavenumber and their position on the Plotly “Greys” colourmap determined the colour of each bar, where the greater the count, the darker was the bar. These were subsequently, overlaid over the averaged spectra for each species, with a standard deviation value being shaded. A common feature for all three species was a strong second derivative significant value at *ca*. 1000 cm^-1^, also identified by the PCA and SOM, attributed to the phenylalanine peak (**Table 2**).

**Table 2.** Mean confusion matrix generated from the PCA-LDA. The values are rounded to the nearest integer.

|  | Precision | Recall | F1-score | Support |
| --- | --- | --- | --- | --- |
| ***C. glutamicum*** | 0.987 | 0.993 | 0.990 | 121 |
| ***M. bovis* BCG** | 0.994 | 0.992 | 0.993 | 118 |
| ***R. erythropolis*** | 0.993 | 0.988 | 0.990 | 121 |
| **Accuracy** |  |  | 0.991 | 360 |
| **Macro average** | 0.991 | 0.991 | 0.991 | 360 |
| **Weighted average** | 0.991 | 0.991 | 0.991 | 360 |

Interestingly, *M. bovis* (BCG) lacks many features shared by the other two species based on the derived barcodes. This was observed previously *via* PCA where *M. Bovis* (BCG) had a different sign for PC1 and *via* the PCA-LDA, where its sign for LD1 was opposite to the others. This indicates that *M. bovis* BCG has a substantially different spectral fingerprint to the other two species analysed, regardless of the method utilised. Discerning between *C. glutamicum* and *R. erythropolis* samples *via* the barcode approach would rely on the lighter shaded barcodes, which correspond to the characteristic traits of each bacterium and their respective spectra.

Overall, standard spectral classification *via* the artificial neural network SKiNET enables high levels of discrimination when analysing multivariate spectral data generated by analysis of bacteria. In contrast, standard PCA is found to exhibit low levels of accuracy. However, this accuracy can be further enhanced using a PCA quadrant analysis, which corroborates the data generated from SOM analysis and SKiNET. Further enhancing the possibility to identify and characterise bacteria is the ability to discern distinct elements of the Raman spectra generated as a result of statistical peak analysis, which affords insight into which components of the spectra yield the greatest significance in terms of determining which peaks have the greatest influence in determining and differentiating the exact organism.

## Conclusions

We have successfully identified and discriminated the Raman spectral fingerprint of the *M. bovis* BCG and two other bacterial genera from the order Mycobacteriales associated with the microbiome of both humans and animals. Despite similarities in physiology between members of the order Mycobacteriales and their ubiquity within the microbiome of both humans and livestock, previous research has not considered the possibility of spectral crossover when utilising Raman spectroscopy. Therefore, the discrimination of these three closely related bacteria provides a valuable insight into the Raman spectra of potentially pathogenic bacteria as well as those within the microbiome. We have shown that a crossover event is highly likely to lead to false positives in the diagnosis of TB using sputum samples especially, if only PCA is utilised. The introduction of the SKiNET algorithm, therefore, affords a higher level of discrimination when compared to PCA alone. By utilising this machine learning technique, we have discriminated between *M. bovis* BCG, *C. glutamicum* and *R. erythropolis* with a high degree of accuracy (*ca*. 99%), which was further enhanced and corroborated using PCA-LDA and PCA quadrant analysis. These methods, all represent a major shift towards spectroscopic diagnosis, which follows a growing trend in industry to move from traditional desktop applications to the Cloud, including office suites, multimedia editing and computer aided design and yet the advantages of connected scalable applications are seldom leveraged in the scientific community.

Whilst the presence of broad peaks can mask certain features of the bacteria (36), and possibly interfere with the detection of *Mycobacterium* sp. when considering the use of biological samples taken from patients, the methods presented herein enable a higher level of discrimination between the bacteria. Future research will consider expansion into the analysis of more Mycobacteriales, which have been readily identified as part of the human microbiome as well as other bacteria, which are also likely to be present within the microbiome as well as the analysis of more pathogenic bacteria as well as various serovars and the analysis of mixed samples, which are more reflective of real-world scenarios. Ultimately, once analysis of these bacteria has taken place individually, a composite culture containing all the respective bacteria would be required and this would enable investigation into the feasibility of Raman as a diagnostic technique for bacterial respiratory illnesses such as TB. Furthermore, the development of individual barcoding for each organism would enable the development of an algorithm which can discriminate between various bacteria and help identify the causative agent of a bacterial respiratory condition. Such an output could be developed into a web-based app that can be accessed globally providing a multi-faceted, qualitative output rather than a quantitative one. In summary, our study provides valuable insights into the combined use of Raman, SKiNET and other multivariate analyses and statistical tools which can rapidly discriminate and identify various bacteria following spectral identification and detection. This enables the important ability to discern the spectra of physiologically similar bacteria especially when compared with more traditional techniques such as PCA along with the ability to generate and consolidate fingerprint barcodes whilst also identifying spectral elements unique to individual bacteria, further adding to the list of tools which are likely to be sought after in the ongoing fight against infectious diseases, especially in a post-Covid world.

## Acknowledgements

We acknowledge funding from the Wellcome Trust (174ISSFPP) and the EPSRC (EP/V029983/1). We thank Dr Jonathan J. S. Rickard, University of Cambridge, for insightful and critical feedback on this study and paper.

## Notes

https://datadryad.org/stash/share/Mv2p7bx7QsOCi3ZzFzASVsyqFOZ5k2oSax07r0aNyKo

### Competing Interest Statement

The authors have declared no competing interest.

